# Human Metalloenzymes Are Disproportionately Linked to Disease: The HEAM Structural Annotation Resource

**DOI:** 10.64898/2026.10.09.758104

**Authors:** Elizabeth H. Miller, Samuel Lim, Matthew Copeland, Joanna S.G. Slusky

## Abstract

Catalytic metal-binding sites are established drug targets for many human diseases, yet systematic study of how metal function relates to disease is limited by the lack of a resource combining structural localization of human metal-binding sites with catalytic annotation at scale. We introduce HEAM (the Human Enzymatic Annotation of Metalloproteins), a structurally annotated dataset of 1,933 human metalloenzymes and 1,171 non-catalytic metalloproteins. HEAM incorporates annotated metals into protein structures at their corresponding binding sites and classifies each site as catalytic or non-catalytic using the machine learning classifier MAHOMES II. HEAM more than quadruples the human site-level coverage of the largest prior metalloenzyme database.

Using HEAM, we show that human metalloproteins are significantly enriched in KEGG disease pathways relative to the human proteome, and that metalloenzymes specifically are significantly enriched in ClinVar pathogenic variants — a distinction obscured in prior, catalytically unresolved metalloprotein datasets. These pathogenic variants are concentrated in metabolic disorders, indicating that metabolic dysfunction is a substantial contributor to metalloenzyme-linked disease. By pairing 3D structural annotation with catalytic classification, HEAM provides both a foundation for structure-guided drug design against metal-binding sites and a resource for determining how catalytic and non-catalytic metal function contribute to human disease.

## Introduction

Metals are ubiquitous in biology and modulate biological function and disease state. In humans, they carry out processes ranging from DNA repair, cell proliferation, and immune regulation including many processes implicated in disease such as cancer metastasis^1^. Metalloenzymes are a particularly important subclass of metalloproteins because their metal-binding sites catalyze many of the reactions in the biological processes listed above. These catalytic metal-binding sites are established drug targets: a range of FDA-approved drugs are small molecules that coordinate the metal in the active site, thereby inhibiting the activity of the metalloenzyme^2^. Knowing whether a metal-binding site is catalytic, along with its structure, is therefore of direct interest to drug development. Catalytic annotation identifies which metal-binding sites are candidates for inhibition, while structural information enables computational analyses that guide the design of molecules to target those sites.

A comprehensive resource containing human metalloproteins, their bound metals, the structural locations of their metal-binding sites, and annotations of whether those sites are catalytic would enable systematic investigation of the roles of metal-mediated function in human biology and disease. Such a resource would also facilitate investigation of how catalytic and non-catalytic metal-binding sites differ in their contributions to disease. However, existing databases each lack some part of the structural and catalytic annotations at the scale needed for systematic analysis.

The Mechanism and Catalytic Site Atlas (M-CSA) contains 1,003 manually curated catalytic sites but does not systematically record bound metals^3,4^. The MetalPDB contains metal-binding sites extracted from PDB structures, but only 50% are functionally annotated, and only a subset of those that are catalytic are annotated as catalytic^5^. The Metal-binding Protein Atlas (MbPA) records metalloproteins with their structures, metal-binding sites, and bound metals, but does not classify which sites are catalytic^6^. The Metalloenzyme Data Bank and Analysis platform (MeDBA) is dedicated specifically to metalloenzymes, but although a subset of its entries were identified by analyzing metal-binding sites in experimentally determined PDB structures, most were identified through keyword searches of UniProt and BRENDA, supplemented by protein family-guided curation from additional databases^7^. The precise structural locations of catalytic metals and their binding sites are therefore unavailable for most entries in the database. With respect to their utility for understanding human biology and disease, the MeDBA is the only one of these databases that provides a straightforward way to filter for human entries.

Recent advances in protein structure prediction and machine learning now make it possible to systematically annotate metal-binding sites and assemble them into a comprehensive resource. Here, we introduce the Human Enzymatic Annotation of Metalloproteins (HEAM), a dataset of human metalloproteins in which each protein has a structure with one or more metals positioned in their binding sites, and each binding site is classified as catalytic or non-catalytic according to whether its metals directly participate in catalysis. HEAM was constructed by retrieving binding site annotations for human proteins from UniProt^8^, retaining those that bind a common biological metal with annotated coordinating residues, computationally placing each metal in the protein’s predicted AlphaFold2^9^ structure at those residues, and classifying each metal-binding site with MAHOMES II, a machine learning classifier that predicts which metal-binding sites are catalytic^10^. To recover additional human metalloproteins lacking metal-coordinating residue annotations in UniProt, HEAM was compared with the MeDBA, and eligible proteins were recovered from experimental metal-bound structures.

Every protein in HEAM includes the location of its metals and metal-binding sites within a structure, together with a catalytic classification for each site. Proteins with at least one catalytic site were classified as metalloenzymes, and proteins with no catalytic metal-binding sites were classified as non-catalytic metalloproteins and were retained for comparison with metalloenzymes in downstream analyses. HEAM contains 1,933 human metalloenzymes and 1,171 non-catalytic metalloproteins. Because every metal-binding site in HEAM has a defined 3D location and catalytic classification, HEAM supports both the identification of catalytic metal-binding sites for structure-guided drug design and the systematic comparison of catalytic and non-catalytic metalloproteins.

To better understand the functional landscape of human metalloproteins, we determined the composition of metal types in HEAM binding sites and the distribution of enzyme classes. We then analyzed the associations of human metalloenzymes and non-catalytic metalloproteins with KEGG biological pathway classes^11–13^, KEGG disease pathways, and ClinVar pathogenic variants^14^, finding that human metalloproteins were significantly enriched in KEGG biological and disease pathways and human metalloenzymes were significantly enriched in ClinVar pathogenic variants.

## Results

### HEAM creation pipeline

The Human Enzymatic Annotation of Metalloproteins (HEAM) dataset was created through five steps—four before comparison with MeDBA and one after (Figure 1).

**Figure 1.**
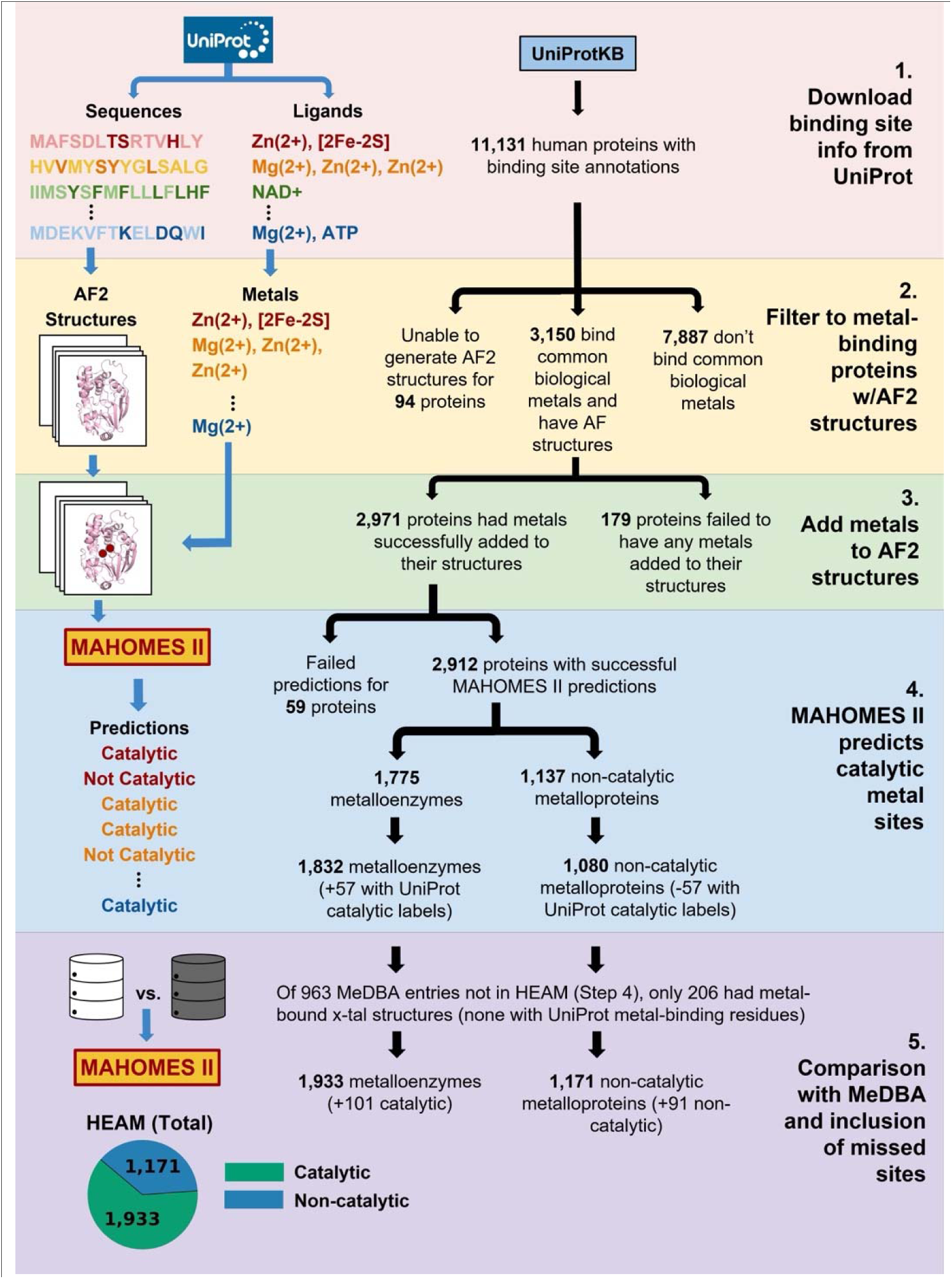
HEAM dataset creation pipeline. Steps 1-5 of the pipeline shown. Step 1—Each UniProt entry has a sequence and ligands annotated with their binding residues (darker colored letters; sequences shown are schematic). Step 2— Proteins that had AF2 structures and common biological metal ligands were identified. Step 3— Metals were placed in the protein’s AlphaFold2 structure via annotated binding residues. Step 4— Each metal-bound structure was input to MAHOMES II, which predicted whether each metal-binding site was catalytic. The resulting dataset, HEAM (Step 4), was compared with the MeDBA. Step 5—Of the MeDBA entries absent from HEAM (Step 4), 206 had metal-bound structures in the PDB. These 206 proteins lacked the UniProt binding site annotations needed to place their metals in AlphaFold structures, so their experimental structures were input to MAHOMES II; 192 returned predictions and 14 failed, and the predictions were added to HEAM, resulting in a total of 1,933 metalloenzymes and 1,171 non-catalytic metalloproteins.

Human proteins carrying UniProt binding site annotations, which record each bound ligand together with the specific residues that coordinate it, were retrieved from UniProtKB. We kept proteins annotated as binding at least one of the common biological metals that MAHOMES II recognizes: Mg, Mn, Fe, Co, Cu, Ni, Zn, and Mo (see Methods for full list of accepted metal ligands, which specifies oxidation states and metal clusters). Although MAHOMES II does not use metal identity as a predictive feature, it considers the atom count of each bound species, which is defined only for these accepted metals; therefore, it can only make predictions for proteins that bind these metal ligands^15^.

Notably, calcium is not included in the list despite being biologically relevant. This is a specific design choice of MAHOMES II: because calcium can often have a structural or regulatory role in an enzyme active site without being part of the catalytic mechanism, it was excluded from the training set of MAHOMES II, and thus calcium was not included in the list of common biological metals that MAHOMES II recognizes.

For each protein sequence bound to a common biological metal, AlphaFold2 (AF2) structures were obtained, and each metal was placed according to the locations of its annotated coordinating residues (see Methods). Most AF2 structures were downloaded directly from the AlphaFold Protein Structure Database (AFDB)^16,17^; the remainder were predicted in-house with AlphaFold2. Pairing curated UniProt binding site annotations with AF2 structures is more reliable than using experimental crystal structures because metals deposited in crystal structures are often of uncertain identity, misannotated^18^, or assigned to uncertain or inaccurate locations^19^. In comparison, UniProt binding site annotations specify both the identity of each metal and the residues that coordinate them, which allows metals to be computationally placed in AF2 structures with correct identities and more accurate locations.

Each annotated metal was added to the protein’s AF2 structure using the “addMetalIon.py” script from the MAHOMES II repository^15^, which places the metal at the centroid of its annotated coordinating residues (see Methods). In MAHOMES II benchmarking against well-resolved experimental structures, this script placed most common biological metals within 1 Å of the resolved metal locations^10^. The resulting metal-bound structures were submitted to MAHOMES II for catalytic site prediction (Fig. 1).

Of the 11,131 human proteins in UniProtKB with binding site annotations, 3,150 both bound a common biological metal and had an AlphaFold2 structure (Fig. 1). The remainder either bound no common biological metal (7,887) or could not be modeled by AlphaFold2 (94) due to unknown residues, excessive length, or prediction failure (see Methods). Metal placement succeeded for 2,971 of the 3,150 proteins, and MAHOMES II returned predictions for 2,912 of these; 179 proteins were removed during metal placement and 59 failed MAHOMES II prediction (see Methods).

Of the 2,912 proteins with successful MAHOMES II predictions, 1,775 (61.0%) had at least one catalytic metal-binding site and were therefore classified as metalloenzymes; 1,137 (39.0%) had none and were classified as non-catalytic metalloproteins (Fig. 1). For the purposes of our dataset labels, a non-catalytic metalloprotein does not necessarily lack enzymatic activity; it just has no catalytic metal-binding sites.

As a midpoint check, we compared the consistency of our enzymatic labeling with UniProt catalytic labeling. UniProt curators annotate some coordinating metals to be catalytic (ligand_note= “catalytic”). Of the 2,912 human metalloproteins in HEAM, only 537 carried a UniProt catalytic metal label (2,375 carried no label on the metal). We interpreted the absence of a UniProt catalytic metal label to be an incomplete annotation rather than evidence that the metal was non-catalytic. Of the 537 UniProt annotated catalytic proteins, MAHOMES II independently predicted 480 (89.4% recall) to be metalloenzymes. The remaining 57 were reclassified as metalloenzymes, changing the composition of metalloenzymes and non-catalytic metalloproteins in HEAM (Step 4) to 1,832 and 1,080, respectively (Fig. 1).

### Comparing HEAM with MeDBA

We compared HEAM to the only other curated metalloprotein dataset with available organismal (and therefore human) annotation, MeDBA, to find any systematic omissions from HEAM and to evaluate the relative successes of the two sets. HEAM and the MeDBA use different definitions of a metalloenzyme. In HEAM, a protein is classified as a metalloenzyme if at least one of its metal-binding sites is catalytic. In contrast, the MeDBA uses a broader definition, including enzymes with metals in catalytic sites (denoted as M-I or M-II, both of which satisfy the HEAM definition of a metalloenzyme), and enzymes in which the metal plays only a structural role in catalysis and is therefore not located in a catalytic site (denoted as M-III)^7^. Of the 1,634 human metalloenzymes in the MeDBA,1,422 are M-I or M-II and 212 are M-III (Fig. 2).

**Figure 2.**
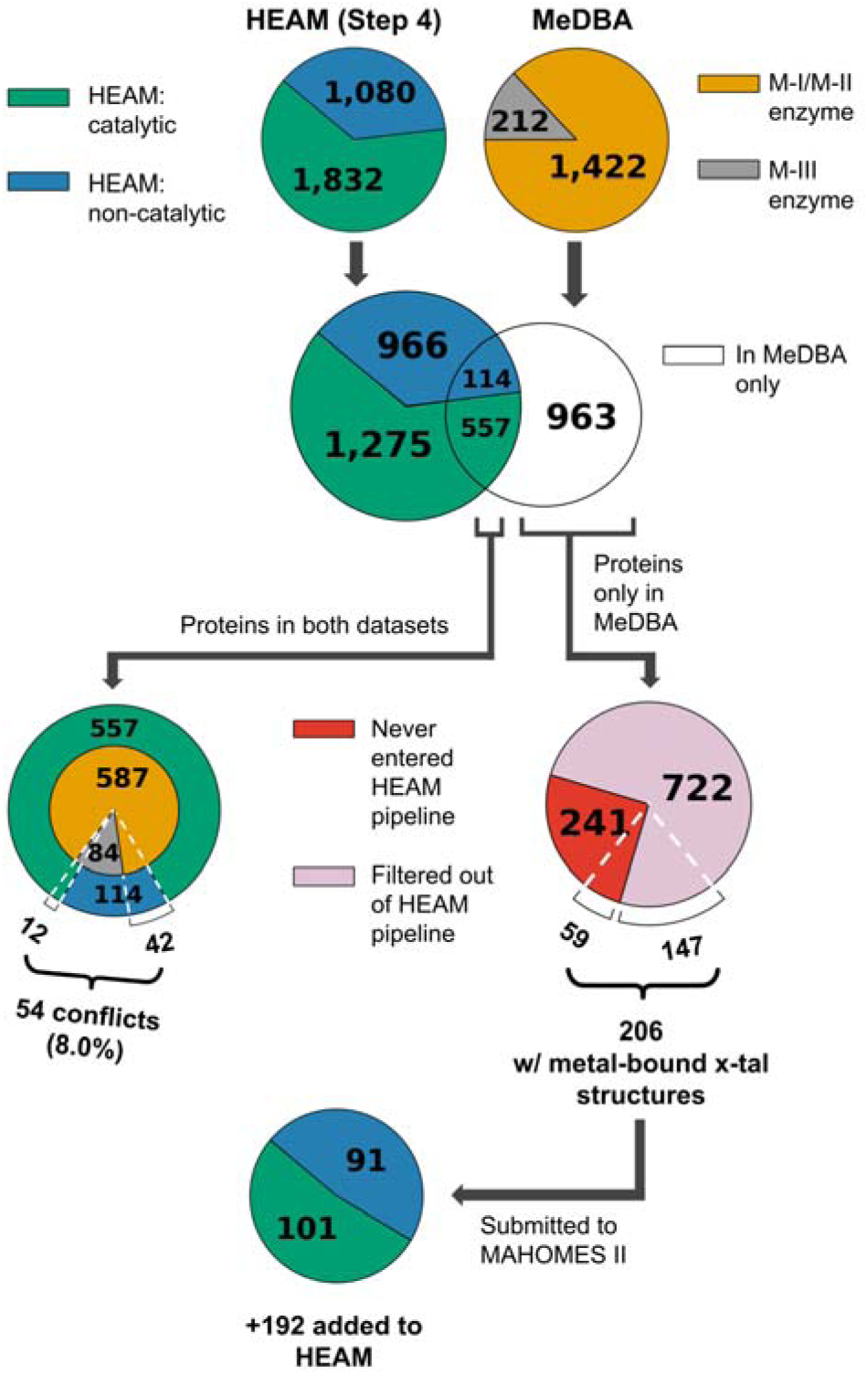
Comparison of MeDBA with metalloproteins in HEAM (Step 4) reveals low overlap. The catalytic composition of each input set is shown at top, with their overlap below. Of the 671 proteins shared by both datasets, 114 were classified as non-catalytic in HEAM (blue). 72 of these proteins are M-III enzymes (gray) in the MeDBA and thus do not represent true disagreements. The true disagreements consist of 42 proteins that are MeDBA M-I/M-II enzymes (yellow) but non-catalytic in HEAM, and 12 proteins that are MeDBA M-III enzymes but catalytic in HEAM (green). c) Of the 963 proteins only in the MeDBA (white), 241 had no UniProt binding site annotations for any ligand (red) and thus could not enter the HEAM creation pipeline; the remaining 722 entered but were removed during filtering (Fig. 1) (pink). Both groups were checked for experimental metal-containing PDB structures; 59 of the 241 and 147 of the 722 (206 in total) had such structures and were submitted to MAHOMES II. Prediction resulted in 101 metalloenzymes and 91 non-catalytic metalloproteins added to HEAM (Step 5).

Only 671 proteins were shared between the 1,634 human metalloenzymes in the MeDBA and the 2,912 metalloproteins in HEAM (Fig. 2). Of these, 617 (92.0%) had agreeing classifications: 545 M-I/M-II enzymes were classified as a metalloenzyme in HEAM, and 72 M-III enzymes were classified as non-catalytic in HEAM (as expected because M-III enzymes fall outside the HEAM definition of a metalloenzyme). The 54 true disagreements (8.0%) consist of 42 M-I/M-II enzymes classified as non-catalytic in HEAM and 12 M-III enzymes classified as a metalloenzyme in HEAM.

### Step 5: Inclusion of MeDBA metalloenzymes absent from HEAM

Of the 963 MeDBA metalloenzymes absent from the HEAM (Step 4), 241 lacked the UniProt binding site annotations required for entry into the HEAM creation pipeline. The remaining 722 had entered the pipeline but were removed during filtering (Fig. 2): 653 were filtered out because none of their annotated ligands corresponded to a common biological metal, and 69 were removed because they either exceeded the 2700-residue limit required for timely AF2 prediction (6), failed metal placement in an AF2 structure (44), or failed MAHOMES II prediction (19). Of these 963 proteins, 128 were M-III.

Worth inclusion in HEAM were 554 of these 653 proteins that have a common biological metal listed as a cofactor in the MeDBA, even though UniProt records no coordinating residues for them. To recover these proteins, each was mapped to experimental PDB structures that already contained the relevant metal cofactors (Methods). Suitable structures were identified for 147 proteins.

The 241 proteins that lacked the UniProt binding site annotations required for entry into the HEAM creation pipeline were similarly checked for metal-bound experimental structures. Suitable structures were identified for 59 of these proteins.

The resulting 206 proteins with metal-bound experimental structures were sent to MAHOMES II, which yielded predictions for 192 proteins, giving 101 metalloenzymes and 91 non-catalytic metalloproteins (Fig. 2). These were added to HEAM, resulting in a total of 1,933 metalloenzymes and 1,171 non-catalytic metalloproteins (n=3,104; Fig. 1). Of these 206 proteins, 60 were M-III.

HEAM is 1.9 times the number of human entries in the MeDBA, containing 1,470 more human metalloproteins. HEAM also has 4 times as many annotated human metal-binding sites as the MeDBA. Among the 671 proteins both in the MeDBA and HEAM before Step 5 of the pipeline, 92.0% had agreeing classifications in both datasets; among the 192 proteins in the MeDBA added to HEAM in Step 5, 79.7% had agreeing classifications (100 M-I/M-II enzymes were classified as catalytic in HEAM and 53 M-III enzymes were classified as non-catalytic in HEAM).

The final size and distribution of HEAM is 3,104 metalloproteins with 3,883 metal-binding sites— 2,124 catalytic and 1,759 non-catalytic.

### Types and distribution of metal-binding sites

Zinc, magnesium, manganese, and iron-binding sites accounted for the majority of metal-binding sites in HEAM. Zinc-binding sites were more frequently non-catalytic than catalytic, consistent with prior predictions of the human proteome containing far more structural than catalytic zinc sites^20^. Magnesium and manganese-binding sites, on the other hand, were more frequently catalytic. Iron-binding sites were likewise predominantly catalytic, in agreement with prior findings that more than half of human iron-binding proteins having a catalytic function^21^. Notably, copper-binding sites were more frequently catalytic in the +2 oxidation state and less frequently catalytic in the +1 oxidation state.

### Enzyme classification

We also determined the frequency of enzyme class within human metalloenzymes. Enzyme classes (EC)^22^ were assigned for 1,931 of the 1,933 HEAM metalloenzymes (Methods). Hydrolases were the largest class (920; 47.6%), followed by transferases (648; 33.6%). 36 metalloenzymes (1.9%) were assigned to more than one enzyme class (darker green, Fig. 3b).

**Figure 3.**
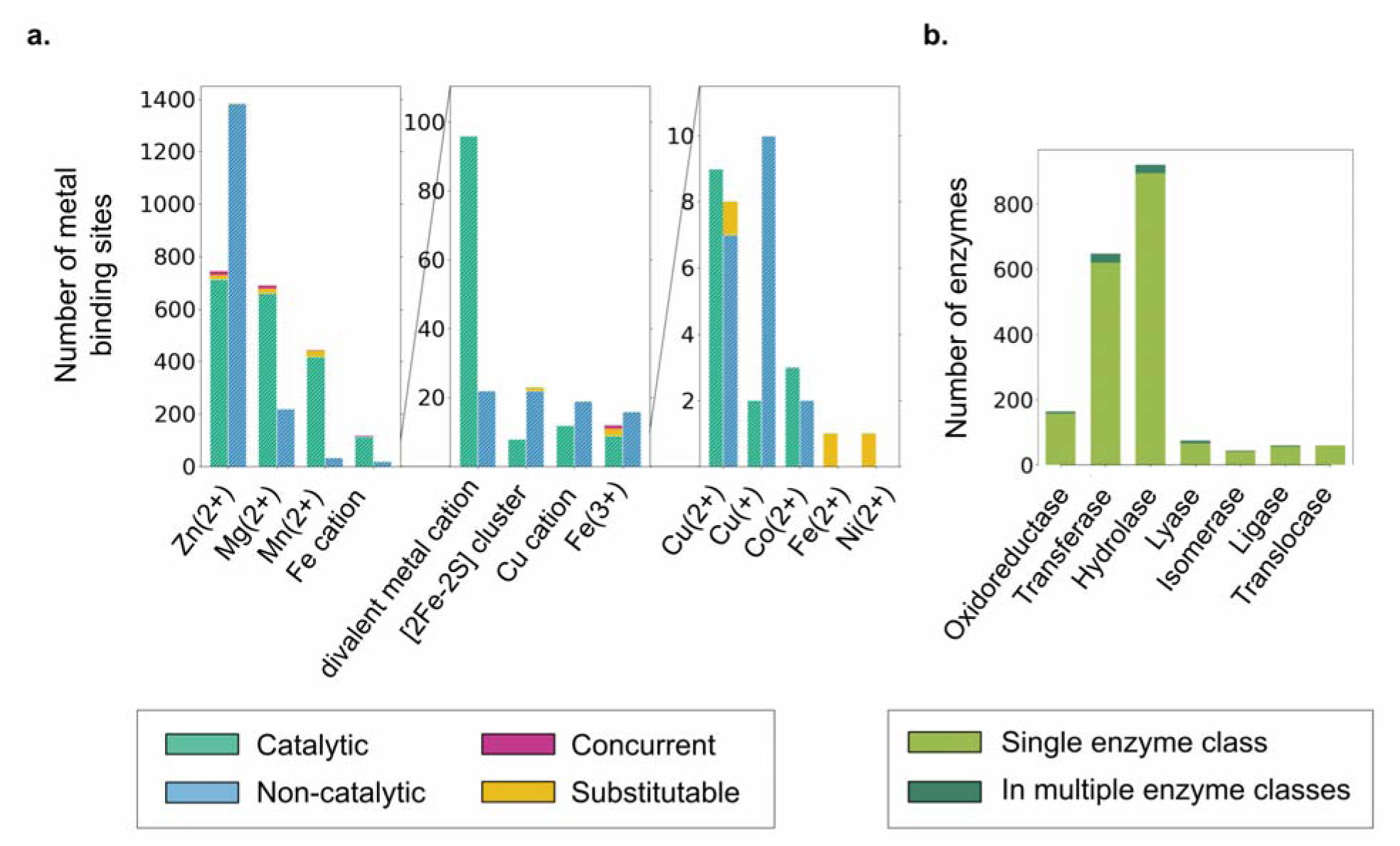
Types and distribution of metal-binding sites across HEAM and enzyme-class assignments of HEAM metalloenzymes. a) Number of sites across HEAM, grouped by metal type, catalytic (green) or non-catalytic (blue). Within each bar, regions are marked by the proportion of sites that bind to other metal types, concurrently (magenta) or via substitution (yellow). The figure is split into three panels with different y-axis scales so that low-count metal sites remain visible. b) Enzyme class distribution across HEAM metalloenzymes (n = 1,931 with an assigned class). Enzymes assigned to more than one class are shown in a darker shade of green. These enzymes are counted once for each class they are assigned.

### Human metalloproteins are significantly enriched in KEGG pathways

Metalloproteins participate in a wide range of biological processes, making their representation across biological pathways an important measure of their functional roles. To determine the relative representation of catalytic and non-catalytic metalloproteins in biological pathways compared to all human proteins, we cross-referenced human proteins to the KEGG pathway database, a curated collection of manually drawn biological pathway maps^11–13^.

To ensure as much biological accuracy as possible, two restrictions were made to the proteins used in this comparison. First, only proteins in the human reference proteome were considered. HEAM proteins absent from the reference proteome were represented by their reference proteome homologs where these existed (Methods), yielding 2,268 metalloproteins— 1,347 metalloenzymes and 921 non-catalytic metalloproteins.

Second, only proteins eligible for KEGG pathway annotation were considered. For a protein to have a gene annotated in a KEGG pathway, the gene must exist in the KEGG GENES database. The KEGG GENES database has a disproportionate fraction of human metalloprotein genes compared to other human proteins genes (human proteins in the reference proteome that aren’t in the 2,268 metalloprotein set used for this comparison): 77.4% of human metalloenzymes and 81.1% of human non-catalytic metalloproteins had genes in the KEGG GENES database, compared to only 12.2% of other human proteins.

Thus, to prevent differences in KEGG GENES coverage between protein groups from influencing the pathway annotation comparison, we restricted the total protein counts in each group to proteins with genes in the KEGG GENES database (catalytic, n=1042; non-catalytic, n=747; other, n=17,787; all, n=19,576). The resulting analysis shows human metalloenzymes and non-catalytic metalloproteins significantly enriched in KEGG biological pathways relative to all human proteins with corresponding genes in the KEGG GENES database (Fig. 4a).

**Figure 4.**
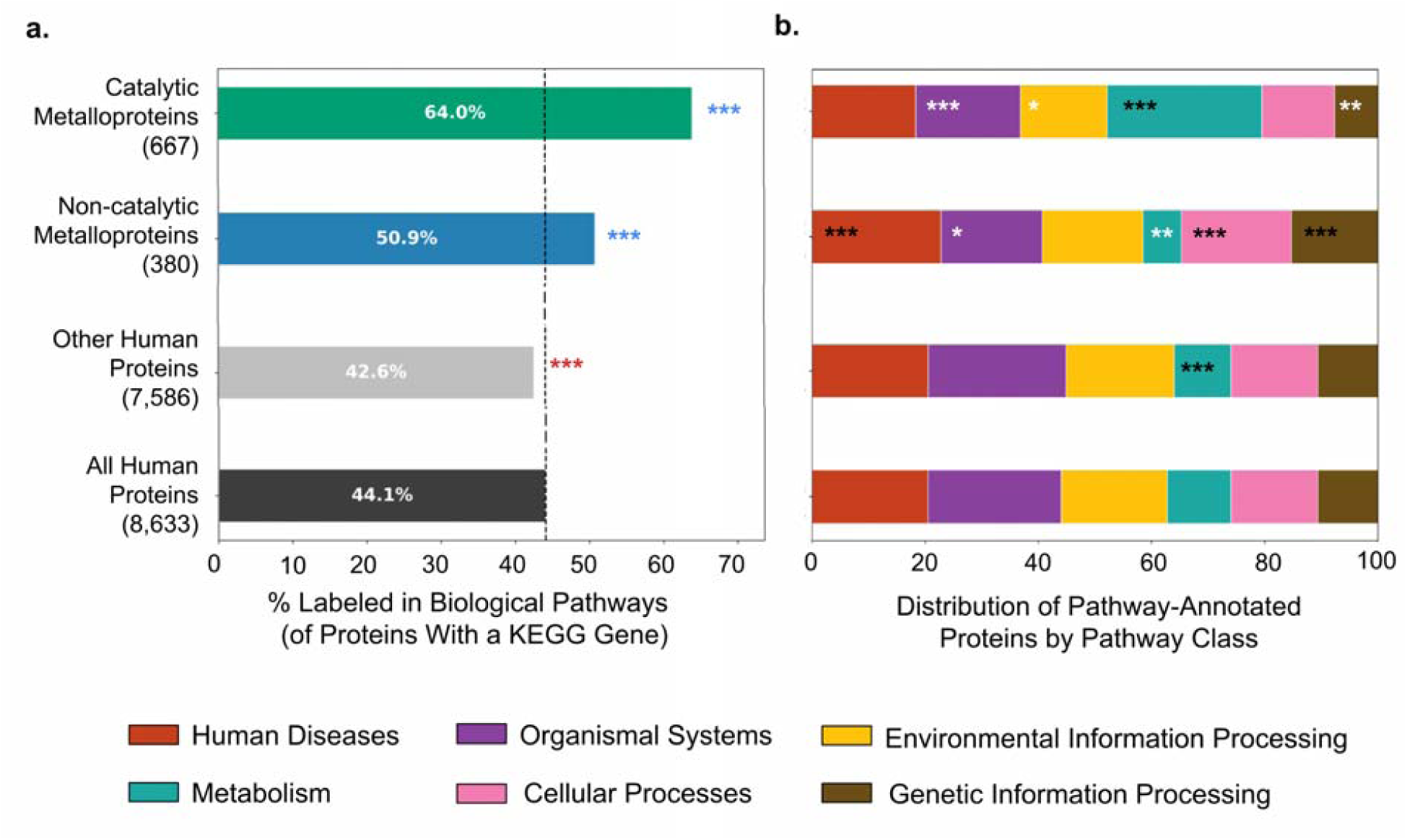
Human metalloproteins are significantly enriched in KEGG biological pathways and distributed differently across pathway classes. a) Percentages of proteins in each group annotated in at least one KEGG biological pathway: human metalloenzymes (green; 667/1,042), human non-catalytic metalloproteins (blue; 380/747), other human proteins (light gray; 7,586/17,787), and all human proteins (dark gray; 8,633/19,576). All four groups are restricted to proteins that appear in the human reference proteome and have a gene in the KEGG GENES database (see Methods). The dashed line marks the percentage of all human proteins annotated in a pathway. Asterisks indicate whether a protein group is significantly enriched (blue) or significantly depleted (red) in biological pathways relative to all human proteins, by binomial test (***, adjusted p < 0.001). b) Distribution of the pathway-annotated proteins in a) across the top-level KEGG pathway classes. For each protein group, each colored segment represents the proportion of pathway-annotated proteins associated with a given top-level KEGG pathway class. A protein is counted once in a pathway class if it has any pathway in that class, and counted separately in each class it spans. Asterisks indicate whether a protein group is significantly enriched (black) or significantly depleted (white) in a given pathway class relative to the fraction of pathway-annotated human proteins in that class, by binomial test (*, adjusted p < 0.05; **, adjusted p < 0.01; ***, adjusted p < 0.001).

#### Representation across KEGG pathway classes

We also compared human metalloenzymes, human non-catalytic metalloproteins, and all other human proteins by the proportion of each whose genes were annotated in each pathway type. KEGG organizes its biological pathway maps into seven top-level classes: Metabolism, Genetic Information Processing, Environmental Information Processing, Cellular Processes, Organismal Systems, Human Diseases, and Drug Development^11–13^. The Human Diseases class consists of 96 disease pathways further analyzed below. To determine how human proteins were distributed among these pathway classes, each protein was counted once for each class in which it had any pathway annotation, regardless of how many pathways within that class it belonged to.

Among the proteins that were annotated in at least one KEGG pathway, human metalloenzymes were significantly enriched in the *metabolism* class, and human non-catalytic metalloproteins were significantly enriched in the *human diseases*, *cellular processes*, and *genetic information processing* classes relative to the fractions of pathway-annotated human proteins in those classes (Fig. 4b). No human protein mapped to a pathway in the *drug development* class, consistent with the fact that this class comprises pathways of drug-related compounds rather than genes^23^.

### Human metalloproteins are significantly enriched in KEGG disease pathways

Beyond their roles in normal biological processes, metalloproteins also contribute to the molecular pathways underlying many human diseases. To determine the relative representation of catalytic and non-catalytic metalloproteins in disease pathways, we cross-referenced human proteins to the KEGG *human diseases* pathway class analyzed above.

Human metalloenzymes and non-catalytic metalloproteins were both significantly enriched in KEGG disease pathways relative to all human proteins with genes in the KEGG GENES database (Fig. 5). 21.9% of human metalloenzymes and 23.3% of non-catalytic metalloproteins were annotated in at least one disease pathway, compared with 15.4% of other human proteins and 16.0% of all human proteins. Non-catalytic metalloproteins were slightly more enriched and showed stronger statistical significance than metalloenzymes (Table S3).

**Figure 5.**
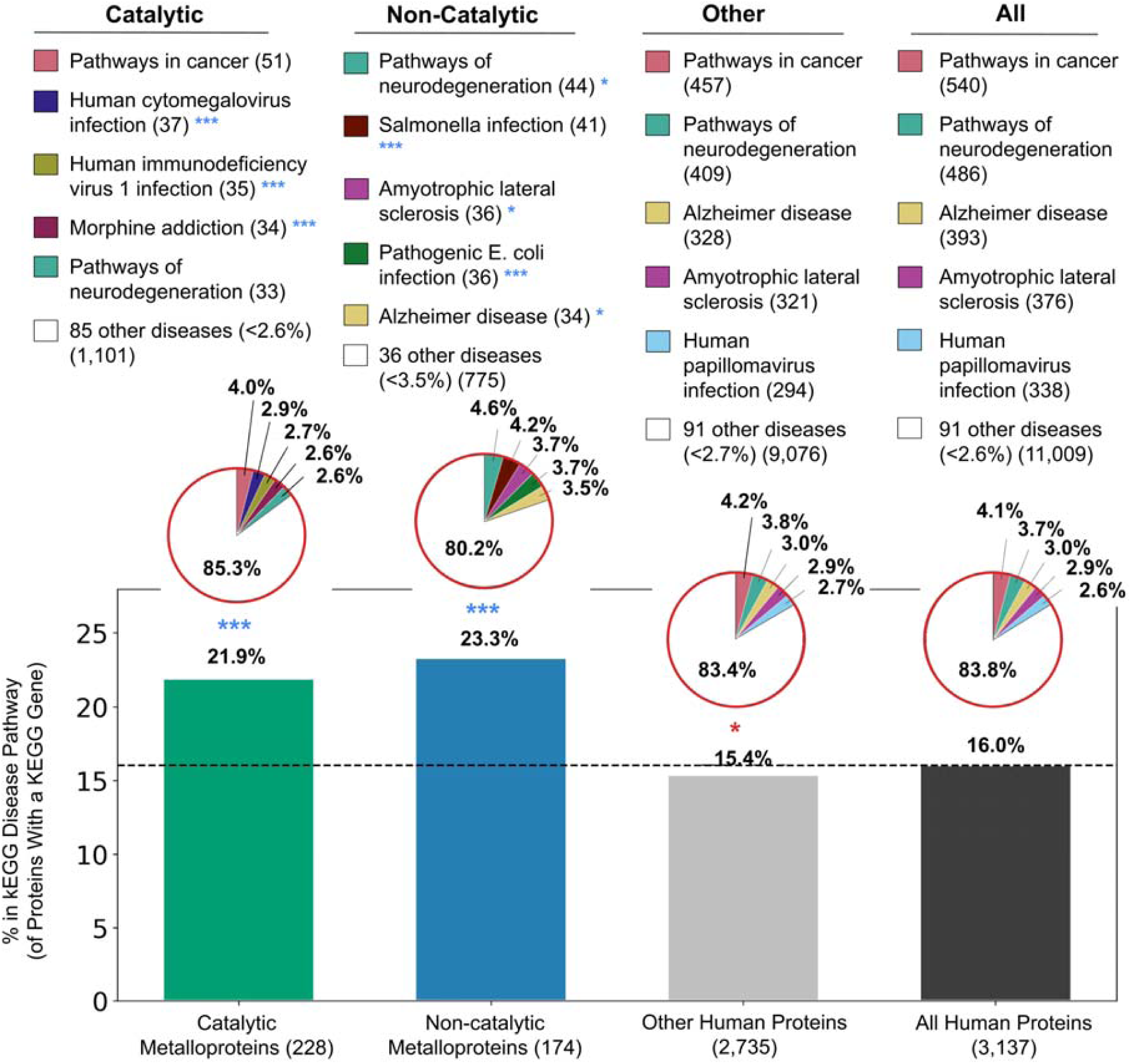
Representation of human protein groups in KEGG disease pathways. The bar graph gives the percentage of proteins in each group whose gene is annotated in at least one human disease pathway: human metalloenzymes (green; 228/1,042), human non-catalytic metalloproteins (blue; 182/747), other human proteins (light gray; 3,726/17,787), and all human proteins (dark gray; 3,137/19,576). All four groups are restricted to proteins that appear in the human reference proteome and have a gene in the KEGG GENES database (see Methods). The dashed line marks the percentage of all human proteins annotated in a disease pathway. Asterisks above the bars indicate whether a protein group is significantly enriched (blue) or significantly depleted (red) in disease pathways relative to all human proteins, by binomial test (*, adjusted p < 0.01; ***, adjusted p < 0.001). The pie chart above each bar shows the distribution of proteins in that group across individual diseases. Because a protein may be associated with more than one disease, proteins can contribute to multiple pie slices. The red outline over each pie indicates that these disease pathways all belong to the KEGG Human Diseases class, shown in red in Figure 4. For each protein group, the legend lists the five diseases with the highest counts of associated proteins; all remaining diseases are grouped as “other diseases”. Asterisks by disease names indicate whether a protein group is significantly enriched (blue) or significantly depleted (red) for a given disease relative to that group’s overall representation among all human proteins annotated in at least one disease pathway, by binomial test (***, adjusted p < 0.001). Full enrichment and depletion results are given in Table S4.

Out of the 96 total disease pathways in KEGG, human metalloenzymes had genes annotated in 90 pathways, and non-catalytic metalloproteins had genes annotated in 86 pathways. Human metalloenzymes were significantly enriched in 20 disease pathways and non-catalytic metalloproteins were significantly enriched in 17, relative to their respective proportions among all human proteins annotated in at least one disease pathway. Human metalloenzymes were significantly depleted in two pathways. Other human proteins had genes annotated in all 96 disease pathways. Of these, they were significantly enriched in 7 and significantly depleted in 29.

Human metalloenzymes were most enriched in morphine addiction, and non-catalytic metalloproteins were most enriched in pathogenic *E. coli* infection (Table S4). Non-catalytic metalloproteins were significantly enriched in several neurodegenerative pathways (Amyotrophic lateral sclerosis, Alzheimer disease, Parkinson disease) (Fig. 5, Table S4).

Among proteins annotated in disease pathways, non-catalytic metalloproteins had a substantially higher proportion of Zn²⁺-binding proteins than metalloenzymes, whereas metalloenzymes had a substantially higher proportion of Mg²⁺-binding proteins (Fig. S1), consistent with our previous analysis (Fig. 3).

### Pathogenic variants show a different distribution

A somewhat different picture of the role of metalloproteins in human disease emerges when comparing proteins with pathogenic variants instead of proteins in disease pathways. ClinVar genetic variant records were analyzed to compare human metalloenzymes, human non-catalytic metalloproteins, and all other human proteins by the proportion of each whose genes carried pathogenic variants. Like the KEGG analyses, proteins were restricted to the human reference proteome, and only proteins with genes in ClinVar were considered (see Methods).

Whereas the proportion of proteins annotated in KEGG disease pathways were significantly greater than that of all human proteins (∼1.4-fold greater for metalloenzymes and 1.5-fold greater for non-catalytic metalloproteins) (Fig. 5), the proportions of each metalloprotein group with pathogenic variants were only ∼1.2-fold greater for metalloenzymes and ∼1.1-fold greater for non-catalytic metalloproteins than that of all human proteins (Fig. 6). Only human metalloenzymes were significantly enriched in pathogenic variants relative to all human proteins.

**Figure 6.**
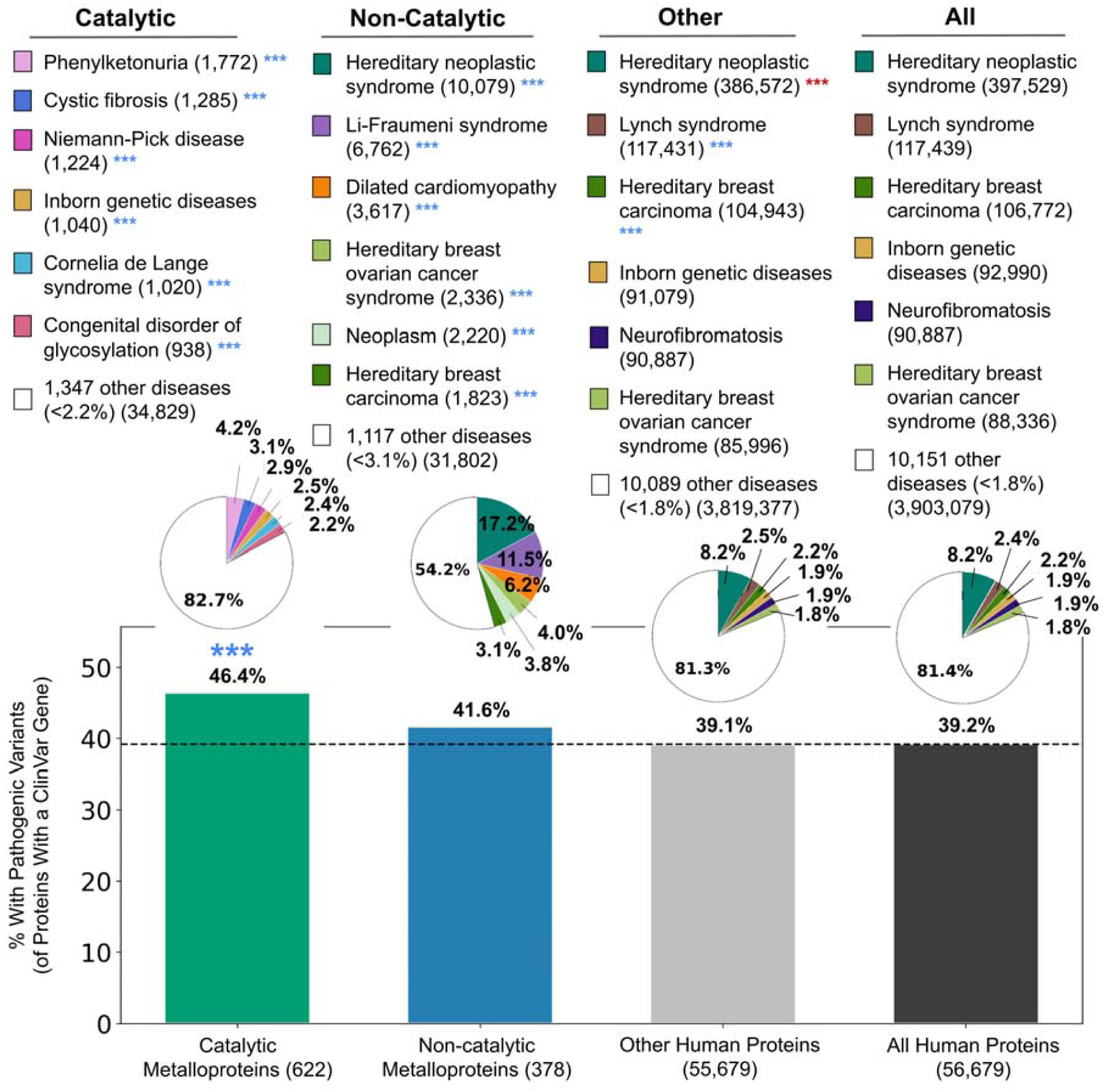
Representation of human protein groups among ClinVar pathogenic variants. The bar graph of the percentage of proteins in each group from genes with at least one genetic variant in ClinVar classified as pathogenic or likely pathogenic: human metalloenzymes (green; 622/1,340), human non-catalytic metalloproteins (blue; 378/908), other human proteins (light gray; 55,679/142,500), and all human proteins (dark gray; 56,679/144,748). All four groups are restricted to proteins that appear in the human reference proteome and have a gene in ClinVar (see Methods). The dashed line shows the 39.2% baseline set by all human proteins. Asterisks above the bars indicate whether a protein group is significantly enriched (blue) or significantly depleted (red) in pathogenic variants relative to all human proteins, by binomial test (***, adjusted p < 0.001). For each protein group, pie charts show the distribution of protein-pathogenic variant associations across disease labels. The legend lists the total number of associations for each disease within each protein group. The six diseases with the highest association counts are shown for each protein group; all remaining diseases are grouped as “other diseases”. Cancers and hereditary cancer predisposition syndromes are in shades of green, and metabolic disorders are in shades of pink. Asterisks by disease names indicates whether a protein group is significantly enriched (blue) or depleted (red) in protein-pathogenic variant associations for a given disease, relative to that protein group’s overall representation among all human protein-pathogenic variant associations, by binomial test (***; adjusted p < 0.001). Full enrichment and depletion results in Table S6.

Across these three groups, pathogenic variants were linked to 17,646 ClinVar diseases, far exceeding the 96 disease pathways in KEGG. To account for the large number of ClinVar diseases with synonymous names, the 17,646 disease labels were clustered down to 10,157 disease labels (see *Methods*). To compare how pathogenic variants were distributed across these diseases, every unique protein-pathogenic variant association was counted once for each disease to which it was linked (Fig. 6). This approach accounts for proteins encoded by genes with multiple pathogenic variants, as well as variants linked to multiple diseases.

For human non-catalytic metalloproteins, four of the six diseases with the greatest numbers of protein-pathogenic variant associations were either a type of cancer or a hereditary cancer predisposition syndrome (green shades, Fig. 6). For human metalloenzymes, three of the seven diseases with the most associations were metabolic disorders (pink shades, Fig. 6).

Each protein group was calculated for statistical enrichment or depletion in diseases relative to that group’s overall representation among all human protein-pathogenic variant associations. Non-catalytic metalloproteins were significantly enriched in each of these four cancer-related diseases, and metalloenzymes were significantly enriched in these three metabolic disorders (Fig. 6, Table S3).

Across all three protein groups combined, hereditary neoplastic syndrome had the greatest number of protein-pathogenic variant associations of any disease; it was significantly enriched in human non-catalytic metalloproteins (Fig. 6) and depleted in both human metalloenzymes and other human proteins (Table S6). Enrichments and depletions for all diseases are recorded in Table S6.

Among proteins with pathogenic variants, non-catalytic metalloproteins had a substantially higher proportion of Zn²⁺-binding proteins than metalloenzymes, whereas metalloenzymes had a substantially higher proportion of Mg²⁺-binding proteins (Fig. S2), consistent with our previous analyses (Figs. 3 and S1).

## Discussion

### HEAM provides the 3D location and catalytic role of human metal-binding sites, over four times the human site-level coverage of the MeDBA

The HEAM dataset occupies a unique niche for drug development because it is a dataset specifically for human metalloproteins. Its defining feature is that it contains a metal-bound structure with annotated catalytic or non-catalytic metal-binding sites for 3,104 proteins— 1,933 metalloenzymes and 1,171 non-catalytic metalloproteins.

Because catalytic metal-binding sites are established therapeutic targets, HEAM’s structures and catalytic/non-catalytic binding site annotations enable downstream structure-guided drug design. By providing the 3D location of each site and labeling which are catalytic, HEAM provides candidate therapeutic target sites in a format directly compatible with structure-guided analyses such as pocket inspection, docking, pharmacophore modeling, and inhibitor design^2^.

By recording metals that bind concurrently within a site and metals that are substitutable with one another, HEAM also provides information about metal occupancy and metal specificity that may be relevant when designing metal-interacting inhibitors.

The most directly comparable resource is the MeDBA, a dataset of metalloenzymes with organismal annotation, which provides structural metal-binding site information based on experimental structures. Although MeDBA uses more reliable structural information than binding sites identified from AlphaFold2 structures, this structural information is only present for 402 of the MeDBA’s 1,634 human metalloenzymes. In total, HEAM contains 3,883 metal-binding sites, compared with 831 metal-binding site annotations in the MeDBA, providing more than four times the site-level structural coverage of human metalloproteins.

### The most commonly occurring metal types are predominantly catalytic, with the exception of Zn^2+^

Across all 3,883 metal-binding sites in HEAM, 55.4% (2,124) are catalytic and 45.9% (1,759) are non-catalytic. This overall predominance of catalytic sites is observed because among the most common biological metal types (Zn^2+^, Mg^2+^, Mn^2+^, and Fe cation), every one except Zn^2+^ is found more in catalytic sites. Because catalytic metal-binding sites are established therapeutic targets, their predominance across HEAM indicates that most human metalloproteins carry a site that may be amenable to therapeutic targeting in this manner.

The fact that Zn^2+^ is not predominantly associated with catalytic sites also illustrates the value of HEAM’s site-level catalytic annotations. Zn²⁺ is known to act both as a catalytic metal, as in carbonic anhydrase and matrix metalloproteinases^2^, and as a structural one, as in zinc-finger domains. Simply knowing that a protein binds Zn²⁺ provides little guidance on what site to target and how to target it. By classifying every Zn²⁺-binding site as catalytic or non-catalytic, HEAM distinguishes catalytically functional Zn²⁺ sites from structural ones, providing the information needed to prioritize sites that are most compatible with established catalytic-site inhibition strategies.

### Human metalloproteins are significantly enriched in KEGG disease pathways, whereas only human metalloenzymes are significantly enriched in ClinVar pathogenic variants relative to all human proteins

To our knowledge, this is the first quantitative comparison of human catalytic and non-catalytic metalloprotein representation across biological pathways, disease pathways, and pathogenic variants, benchmarked against the overall representation of the human proteome.

Human metalloproteins are significantly enriched in KEGG disease pathways relative to all human proteins of the reference proteome that have a gene in the KEGG GENES database. The percentage of metalloenzymes annotated in disease pathways is ∼1.4-fold greater than that of all human proteins, and the percentage of non-catalytic metalloproteins is ∼1.5-fold greater.

However, the percentage of metalloenzymes with pathogenic variants in ClinVar is only ∼1.2-fold greater than that of all human proteins, and the percentage of non-catalytic proteins is only ∼1.1-fold greater. Additionally, only metalloenzymes are significantly enriched in pathogenic variants relative to all human proteins.

The greater relative difference in annotation percentages between human metalloproteins and all human proteins for KEGG disease pathways (∼40-50% greater) than for ClinVar pathogenic variants (∼10-20% greater) may be due to the fact that KEGG disease pathways include genes that are mutated in disease as well as genes whose normal functions are implicated in disease processes. In contrast, ClinVar only catalogs genes with documented pathogenic variants^22^. Thus, the greater KEGG difference may reflect its broader annotation criteria.

Although non-catalytic metalloproteins have a higher percentage of proteins in KEGG disease pathways than metalloenzymes (23.3% vs. 21.9%), they have a lower representation compared to metalloenzymes among proteins with ClinVar pathogenic variants (41.6% vs. 46.4%). This may be because pathogenic variants of metalloenzymes disproportionately cause disease through disruption of metabolic functions.

Human metalloenzymes are significantly enriched in the KEGG *metabolism pathway* class, whereas non-catalytic metalloproteins are significantly depleted in it relative to the percentage of pathway-annotated proteins in that class (Fig. 4b). Additionally, three of the seven diseases with the highest metalloenzyme-pathogenic variant counts are metabolic disorders (phenylketonuria, Niemann-Pick disease, congenital disorder of glycosylation), none of which are represented among the KEGG disease pathways analyzed. Metalloenzymes are significantly enriched in each of these diseases relative to their representation among all human protein-pathogenic variant associations (Fig. 6), while non-catalytic metalloproteins are significantly depleted in both phenylketonuria and Niemann-Pick disease (Table S6). Together, these findings support that disruption of metabolic functions may contribute to metalloenzymes having a higher percentage of proteins with pathogenic variants than non-catalytic metalloproteins.

### Comparison with previous computational metalloprotein annotations

Previous computational studies have estimated the number of human proteins that bind Zn^2+^, Mg^2+^, and inorganic iron.

Andreini et al.^24^ estimated that roughly 2,800 human proteins bind Zn^2+^ in vivo, combining three approaches: X-ray Zn^2+^-binding patterns, Pfam metal-binding domain libraries, and human gene sequence annotations^24^. HEAM contains only 1,660 Zn^2+^-binding proteins in comparison (59%). This discrepancy is mostly due to the limited coverage of UniProt binding-site annotations, which identify only 1,671 Zn^2+^-binding human proteins; the HEAM creation pipeline contributes only a minimal additional reduction.

Piovesan et al. assigned putative Mg^2+^-binding sites to 3,751 human sequences^25^, while HEAM contains only 848 Mg^2+^-binding proteins (23%). This larger discrepancy is likely partly due to lack of UniProt binding-site annotations and partly due to the mechanism of annotation used by Piovesan et al. Specifically, Mg^2+^-binding sites were transferred from experimentally resolved templates to homologous sequences, including templates from other organisms and below 30% sequence identity, likely leading to over labeling.

However, HEAM recovers more inorganic iron-bound proteins than Andreini et al.^21^ predicted. Andreini et al. used iron-binding Pfam domains and structure-derived coordination patterns to predict that approximately 2% of human genes encode iron-binding proteins, including 139 genes encoding proteins that bind inorganic iron ions^21^. In contrast, HEAM contains 157 genes that bind to individual iron ions, 13% more than the Andreini et al. estimate.

Together, these comparisons indicate while HEAM expands on previous annotation and coverage of human metal-binding sites, and it contains more inorganic iron-bound proteins than previous estimates, it remains a conservative representation compared to previous computational estimates of Zn^2+^-binding and Mg^2+^-binding human proteins. This conservative bias makes HEAM reliable for metal-binding annotation, while also points to exciting future opportunities to expand HEAM’s coverage as additional true metalloproteins receive experimental annotation. It will be important to continue to monitor the relationship between metalloproteins and human diseases through further annotation expansion.

## Methods

### HEAM creation pipeline

#### i. Protein set retrieval and identification of common biological metal-binding proteins

Human proteins carrying UniProt binding site annotations were retrieved from UniProtKB (release 2025_03, June 2025). These annotations record each bound ligand together with the specific residues that coordinate it. Both reviewed and unreviewed entries were included using the query "(taxonomy_id:9606) AND (ft_binding:*)", returning 11,131 proteins. For every protein, the annotations were parsed to record each bound ligand, the residues coordinating that ligand, and the evidence source(s) supporting each coordinating residue.

The UniProt binding site annotations were then used to identify proteins binding any ligands classified as common biological metals. The ligands treated as common biological metals in this paper are the metals that MAHOMES II is programmed to recognize: Mo, MoO_4_^2-^, Mo^4+^, Mo^6+^, dioxothiomolybdenum (VI) ion, Ni^3+^, Ni^2+^, Zn^2+^, MgF_3_^-^, Mn^3+^, Mn^2+^, Co^2+^, FeO, Fe^2+^, Fe_2_O, Fe^3+^, [2Fe-2S] cluster, Cu^2+^, Cu_2_O, Cu_2_, Cu^+^, and Co^3+^. MAHOMES II considers the atom count of each bound species as a predictive feature, and atom counts are defined only for these accepted metal ligands; thus, proteins binding other metal ligands cannot be predicted by the model^15^.

#### ii. Structure retrieval

Structural models were obtained for proteins identified as binding a common biological metal as follows: where available, the AlphaFold2 (AF2) structure was downloaded from the AlphaFold Protein Structure Database (AFDB) in PDB format. Proteins without an AFDB structure were predicted with AlphaFold2. Sequences containing selenocysteine, which AlphaFold2 does not support^17^, had those residues substituted with cysteine prior to prediction.

Structures could not be obtained for 94 of the 3,244 proteins that bound a common biological metal. 93 of these were excluded from AlphaFold2 prediction because they either contained unknown residues (68) or had more than 2700 residues (25). The 2700-residue limit matches the length cutoff used for the construction of the AlphaFold human proteome dataset, beyond which predictions become computationally impractical due to memory and runtime constraints^9^. The remaining protein was submitted for prediction but failed due to HHBlits.

#### iii. Metal ion placement

For each protein with an AF2 structure, the parsed annotations were used to determine every occurrence of a common biological metal and its coordinating residues. Of the common biological metals listed above, Ni^2+^, Zn^2+^, Mn^2+^, Co^2+^, Fe^2+^, Fe^3+^, [2Fe-2S] cluster, Cu^2+^, and Cu^+^ were identified in these annotations, and each occurrence of these metal ligands was added to the protein’s structure using the “addMetalIon.py” script from the MAHOMES II repository^15^. This script performs metal placement only and is independent of the MAHOMES II predictive model.

Ligands in UniProt annotated to be “a divalent metal cation”, “Fe cation”, or “Cu cation” were also considered to represent common biological metals. To add these metals to their proteins’ AF2 PDB files, the PDB metal codes ZN, FE, and CU, respectively, were used to represent them. Because MAHOMES II uses the atom count of the bound species as a predictive feature rather than metal identity, the specific metal identity used does not affect the prediction, provided that the selected metal is among those encoded for by the model.

Because UniProt reports only the sequence positions of the coordinating residues, “addMetalIon.py” first parsed the protein’s PDB file to determine each coordinating residue’s identity. For each residue type the script encodes, which we extended to include tryptophan and tyrosine, it identified the sidechain atom most likely to coordinate a metal. The metal was placed at the centroid of each protein’s coordinating atoms, and the metal and its coordinates were written to the PDB file. A metal was omitted if none of its coordinating residues were among the residue types encoded for by the script, or if any two of its coordinating residues were more than 12 Å apart, which was taken to indicate that they did not form a single binding site.

This method worked for every common biological metal that appeared in the UniProt data except for the [2Fe-2S] cluster. For this ligand, a slightly different method was used to add it to the protein’s PDB file: since it was a cluster and not a metal ion, “addMetalIon.py” was used to find the coordinates of the cluster centroid. The coordinates of the two iron and two sulfur atoms were derived by arranging the atoms as a planar, symmetric rhombus centered on the cluster centroid, using an Fe-Fe bond distance of 2.70 Å and an Fe-S bond distance of 2.25 Å.

#### iv. Separating metals inferred to bind at different times

UniProt annotates every metal reported to bind a protein regardless of whether those metals are present simultaneously, so a single PDB file could contain metals that are never physically bound at the same time. Submitting such a file to MAHOMES II unchanged would yield unreliable predictions, because although MAHOMES II does not consider metal identity, it does consider the number and locations of coordinating metals in a site^10^. Each PDB file containing metals inferred to bind at different times was therefore replaced by a set of derived PDB files, all with the same protein coordinates and each containing only metals inferred to bind concurrently.

Metals were classified as binding at different times if they shared identical coordinates, or they were within 5 Å of one another and shared no supporting UniProt evidence source. Metals were then partitioned into groups such that every pairwise combination of metals within a group was classified as binding concurrently; each group was written to its own derived PDB file, retaining the original protein coordinates but containing only that group’s metals.

#### v. MAHOMES II prediction

All resulting PDB files were submitted to MAHOMES II on the University of Kansas high-performance computing cluster for catalytic site prediction. Of the 59 failures, 46 arose from MAHOMES II-specific errors, 12 from memory limits, and 1 from a missing Bluues^26^ calculation.

For proteins with multiple derived PDB files, MAHOMES II predictions for a given site were consistent across all files, even when the number of metals differed. Each protein and each unique site was therefore counted only once in all downstream analyses.

#### vi. Reclassification using UniProt catalytic metal annotations

Ligands in UniProt binding site annotations are sometimes annotated as catalytic (ligand_note=“catalytic”). These annotations were used to identify false negatives among the MAHOMES II predictions. Every protein predicted to be non-catalytic was checked against these annotations, and such proteins with at least one catalytic annotation were reclassified as metalloenzymes, moving 57 proteins from the non-catalytic set to the metalloenzyme set.

### Supplementing HEAM: Recovery of Additional Human Metalloproteins via the MeDBA

Human metalloenzyme entries were downloaded from the MeDBA (January 2026), a curated database of metalloenzymes assembled from experimental PDB structures, keyword searches of UniProt and BRENDA, and other databases. These entries were cross-referenced with the metalloproteins in HEAM (Step 4) to compare the size and classifications of the two datasets and to identify metalloenzymes the HEAM dataset construction pipeline may have missed.

Of the 1,634 MeDBA human metalloenzymes, 963 proteins were absent from HEAM (Step 4) and were checked against the parsed annotations from the initial UniProtKB download (release 2025_03, June 2025). 241 had never entered the HEAM creation pipeline because they lacked UniProt binding-site annotations, and 722 had entered the pipeline but were subsequently filtered out.

Among the 722 filtered proteins, 653 were removed because none of their UniProt-annotated ligands was a common biological metal. However, of these 653, 554 had a common biological metal listed as a cofactor in the MeDBA and were therefore considered for recovery. Similarly, of the 241 proteins that never entered pipeline, 207 had a common biological metal listed as a cofactor in the MeDBA and were considered for recovery. Together, these two groups gave 761 candidate proteins for recovery.

Because these proteins lacked the residue-level annotations required to place metals in AlphaFold2 structures, experimentally determined structures were used, as these already contained the bound metals. Each selected protein was mapped from its UniProtKB accession number to corresponding RCSB PDB entries using the UniProt ID-mapping service (UniProtKB → PDB) (February 2026). 537 proteins had at least one mapped structure; the remaining 224 had none. The mapped structures were downloaded from the RCSB PDB in mmCIF format and converted to PDB format using GEMMI v0.6.5^27^.

Because a single UniProt accession number can map to multiple experimental structures, and these structures are not always monomers, one representative chain was chosen per protein. Candidate chains were drawn from all chains across the mapped structures, keeping only those that contained the complete set of metal cofactors listed for the protein in the MeDBA. Each candidate chain was aligned to the canonical UniProt sequence using the PairwiseAligner from Biopython v1.84^28^ with the default settings. The alignment score was divided by the chain length and, separately, by the UniProt sequence length. The product of these two ratios was used as a composite alignment score to rank candidate chains, favoring chains that aligned well over their own length while covering most of the UniProt sequence. The chain with the highest product was selected.

Of the 537 proteins with experimental PDB structures, 331 were excluded because none of their structures contained a chain with the complete set of metal cofactors listed for that protein in the MeDBA. The selected chains for the remaining 206 proteins were submitted to MAHOMES II. Predictions succeeded for 192 proteins, which were subsequently added to HEAM.

### Enzyme Classification

Enzyme Commission (EC) numbers were assigned to HEAM metalloenzymes from two sources: existing UniProt annotations and, where these were absent, predictions from CLEAN, a machine learning algorithm that predicts EC numbers from protein sequences^29^. Each metalloenzyme was checked for a UniProt EC annotation (release 2026_01, May 2026); 1,387 had one and were assigned it, and the rest were classified with CLEAN using the CLEAN web server (https://clean.platform.ibiofoundry.illinois.edu).

CLEAN accepts single sequences of at most 1,022 residues. The 514 proteins at or below this length were submitted directly. For the 13 proteins exceeding 1,022 residues, the domain containing the metal binding site was identified in PyMOL v.3.1.0^30^, and its sequence (< 1,022 residues) was submitted to CLEAN in place of the full-length sequence. 2 of these 13 proteins were not submitted to CLEAN because their metal site domains were discontinuous in sequence. These 2 proteins remain in HEAM without an EC assignment, giving 1,931 metalloenzymes with an assigned enzyme class.

EC numbers were used only to assign each metalloenzyme to a top-level enzyme class. Where a metalloenzyme carried multiple EC numbers falling within a single top-level class, it was counted once in that class. Where its EC numbers spanned more than one top-level class, it was counted once in each class it spans (Fig. 3b).

### KEGG Pathway Analysis

#### i. Construction of protein sets for pathway comparison

For this comparison, human proteins were defined as the entries in the UniProtKB human reference proteome (UP000005640, release 2026_02), comprising 147,506 entries including isoforms arising from alternative splicing.

HEAM was constructed by filtering from a UniProtKB query of all human entries with binding-site annotations. Because this query was not restricted to the reference proteome, some HEAM proteins have no reference proteome entry. To enable comparison against the reference proteome, the metalloprotein set used for this KEGG pathway analysis comprised HEAM proteins present in the reference proteome together with reference proteome homologs of those absent from it.

Of the 3,104 metalloproteins in HEAM, 2,148 were present in the reference proteome. The remaining 956 proteins were aligned to the reference proteome using BLASTp v.2.17.0. A protein sequence was considered to have a reference proteome homolog if it had greater than 90% sequence identity to a protein in the reference proteome. Of the 956 HEAM proteins not in the reference proteome, 788 were homologous to one of the 2,148 HEAM proteins already present in the reference proteome, 147 proteins were homologous to 120 non-HEAM proteins in the reference proteome, and 21 were not homologous to any protein in the reference proteome.

The 2,148 HEAM proteins in the reference proteome and the 120 reference proteome homologs of HEAM proteins not in the reference proteome were used as the human metalloprotein set for this KEGG pathway comparison. Catalytic and non-catalytic classifications for the 120 non-HEAM proteins were assigned based on the classifications of their HEAM homologs. This resulted in a total of 2,268 human metalloproteins (1,347 catalytic, 921 non-catalytic).

These human metalloproteins, along with the remaining 145,238 proteins in the reference proteome, were further filtered down to only proteins that had a gene in the KEGG GENES database, resulting in 1,042 metalloenzymes, 747 non-catalytic metalloproteins, and 17,787 other human proteins, for a total of 19,576 human proteins.

#### ii. Biological pathway mapping

The 19,576 human proteins with genes in the KEGG GENES database were mapped to KEGG pathways using the KEGG REST API. UniProtKB accession numbers were converted to human KEGG gene identifiers, each of which was linked to its corresponding set of associated KEGG pathways.

Each mapped pathway was assigned to one of the seven top-level KEGG pathway classes using the KEGG BRITE hierarchy. Disease pathways were defined as the 96 pathways constituting the KEGG Human Diseases pathway class.

#### iii. Statistical analysis

Enrichment and depletion of KEGG pathway annotation among human metalloenzymes, non-catalytic metalloproteins, and other human proteins were assessed using two-sided binomial tests, each comparing an observed number of successes in *k* in *n* trials against a background proportion *p*. The expected count was *n* × *p*, and fold change was calculated as the observed count divided by the expected count. P-values were adjusted using the Benjamini-Hochberg procedure^31^, and tests significant after adjustment (adjusted p < 0.05) were classified as enriched (fold change > 1) or depleted (fold change < 1), graded at three thresholds (adjusted p < 0.05*, < 0.01**, and < 0.001***). Four tests were performed:

1. For the biological pathway annotation rate (Fig. 4a), successes were the group’s proteins annotated in at least one biological pathway, trials were the group’s total number of proteins, and the background proportion was the annotation rate across all human proteins with a gene in the KEGG GENES database.
2. For the disease pathway annotation rate (Fig. 5 bar plot), the same quantities for 1) were used but restricted to the 96 disease pathways
3. For the pathway class distribution (Fig. 4b), the test was restricted to proteins annotated in at least one biological pathway: successes were the group’s annotated proteins appearing in the pathway class, trials were the group’s total number of annotated proteins, and the background proportion was the class’s share of all annotated human proteins
4. For individual disease pathways (Fig. 5 pie plot), the test was restricted to proteins annotated in at least one disease pathway: successes were the disease pathway’s proteins belonging to the group, trials were the total number of proteins annotated in that pathway, and the background proportion was the group’s share of all annotated human proteins

### ClinVar Pathogenic Variant Analysis

To identify human metalloproteins whose genes carry pathogenic variants linked to diseases, “variant_summary.txt”, a tab-delimited bulk summary of all ClinVar variant records, was downloaded from ClinVar (June 2026).

Human proteins were mapped to ClinVar entries using the primary gene symbol recorded in each protein’s UniProt entry. ClinVar entries for these genes were retained if they used the GRCh38 assembly and were annotated with a clinical significance of “Pathogenic” or “Likely pathogenic”. As in the KEGG pathway analysis, proteins were restricted to the human reference proteome and divided into the same three groups: metalloenzymes, non-catalytic metalloproteins, and other human proteins. Because a protein can carry pathogenic variants only if its gene is present in ClinVar, all analyses were further restricted to proteins whose gene symbol matched a ClinVar record, resulting in 1,340 metalloenzymes, 908 non-catalytic metalloproteins, and 142,500 other human proteins, for a total of 144,748 human proteins.

Synonymous disease names were consolidated as described below. Two comparisons were then made across human metalloenzymes, non-catalytic metalloproteins, and other human proteins. The first tested each group for enrichment or depletion in the proportion of its proteins carrying pathogenic variants, relative to all human proteins with a gene in ClinVar. The second tested each group within individual diseases linked to human pathogenic variants, relative to that group’s overall share of all protein-pathogenic variant associations.

#### i. MONDO-based disease name consolidation

In ClinVar, a single disease can be represented by multiple names (e.g., “Lynch syndrome” is also known as “Hereditary nonpolyposis colon cancer” and “Hereditary nonpolyposis colorectal carcinoma”). Each pathogenic variant entry may be linked to one or more disease names, each carrying phenotype identifiers from one or more resources within the ClinVar dataset (e.g., MONDO^32^, Orphanet^33^). To reduce redundancy, disease names were consolidated using MONDO disease identifiers. MONDO organizes diseases into a hierarchical ontology in which more specific disease names are the children of broader parent names, and each name is associated with recognized synonyms.

For each disease name with a MONDO identifier, that identifier was used, and other identifiers listed for the same disease name were not considered further. For each disease name without a MONDO identifier, the MONDO ontology file, “mondo.obo”, (June 2026) was searched to determine whether the disease name was listed as a synonym or as an obsolete term mapped to a current MONDO identifier. If so, the corresponding MONDO identifier was used; otherwise, the disease name was retained under its original source identifier(s).

For each pathogenic variant, the set of MONDO identifiers collected across its associated disease name entries was reduced to the most specific disease names using the MONDO ontology graph: any MONDO identifier that was an ancestor of another identifier in the same set was discarded, retaining only the most specific terms. This was done using obonet v.1.3.0^34^ and NetworkX v.3.2.1^35^ to parse and traverse the MONDO graph.

Following MONDO-based disease name consolidation, each disease name and the number of associated protein-pathogenic variant pairs were recorded for further consolidation of remaining synonymous disease names. Protein-pathogenic variant pairs, rather than variants alone, were counted because a single gene can correspond to multiple distinct proteins in the dataset. This consolidation condensed 17,646 disease names to 12,905 disease names.

#### ii. AI-assisted disease name consolidation

The MONDO identifier-based matching did not resolve all disease names referring to the same conditions. Claude Code v.2.1.116^36^ was used to handle the remaining cases. The model was given the 12,957 disease names produced by the MONDO-based consolidation, each with its count of associated protein-pathogenic variant pairs, and instructed to performe three operations (full prompt in Supplementary Text S1):

(a) identify entries combining several distinct diseases under a single name (e.g., “Marfan Syndrome/Loeys-Dietz Syndrome/Familial Thoracic Aortic Aneurysms and Dissections”) and split them into individual disease names;
(b) merge disease names that are synonyms of one another;
(c) merge disease names describing the same phenotypic condition arising from different genes (e.g., “Joubert syndrome 38” and “Joubert syndrome 39”, merged into “Joubert syndrome”).

Protein-pathogenic variant pair counts were updated after each operation. Merged entries were assigned the sum of their component counts; for split entries, the component count was assigned to each.

The model recorded the disease names involved in each operation and a justification for it. Because model output is not deterministic, the complete record of these operations and their justifications is provided in Table S7, allowing each decision to be assessed directly.

This consolidation condensed 12,905 disease names to 10,157, comprising 1,019 splits, 111 merges of synonymous names, and 2,753 merges of names sharing a phenotype.

#### iii. Enrichment testing

Enrichment and depletion of ClinVar pathogenic variant annotation among human metalloenzymes, non-catalytic metalloproteins, and other human proteins were assessed using two-sided binomial tests, following the same conventions as the KEGG pathway analysis.

Two tests were performed:

1. For the pathogenic variant rate (Fig. 6 bar plot), successes were the group’s proteins carrying at least one pathogenic variant, trials were the total number of proteins in the group, and the background proportion as the rate across all human proteins with a gene in ClinVar
2. For individual diseases (Fig. 6 pie plot), successes were the disease’s protein-pathogenic variant associations belonging to the group, trials were the total number of associations linked to that disease, and the background proportion was the group’s share of all protein-pathogenic variant associations

## Data availability

The HEAM database is freely available at https://heam.ku.edu/. For each human protein, HEAM provides a metal-bound structure with three-dimensionally located metal-binding sites, each classified as catalytic or non-catalytic using the MAHOMES II classifier. Users can search and filter entries through the web interface (https://heam.ku.edu/search) or download the complete database (https://heam.ku.edu/download/all). The database and filtered search results are distributed under a Creative Commons Attribution-NonCommercial-ShareAlike 4.0 International (CC BY-NC-SA 4.0) license.

## Code availability

The command-line pipeline used to generate the HEAM database and generate the metal-bound structures for each protein can be found at https://github.com/SluskyLab/heam-pipeline.

## Acknowledgements

We gratefully acknowledge helpful edits from members of the Slusky lab. We also appreciate funding for this project from the KU Endowment.

