## supplemental info for "Human Metalloenzymes Are Disproportionately Linked to Disease: The HEAM Structural Annotation Resource"

**Supplementary Data**

**Figure S1. Distribution of KEGG disease pathway-annotated human metalloproteins by metal type.** The bars show human metalloenzymes (n = 228) and non-catalytic metalloproteins (n = 174) annotated in at least one KEGG disease pathway and present in the UniProt reference proteome. Each colored segment within a bar represents the proportion of disease pathway-annotated proteins that bind a given metal type. A protein is counted once for each metal type it binds.
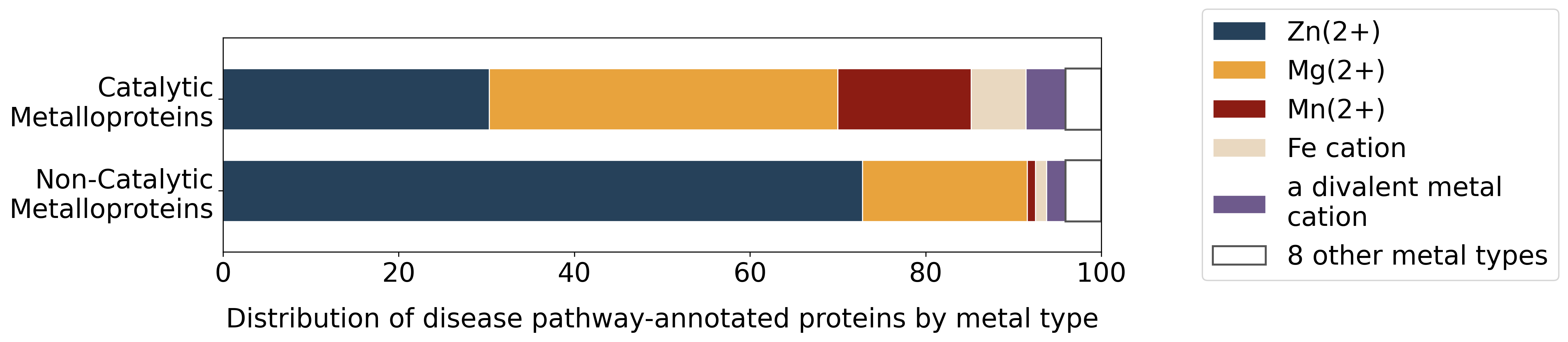

**Figure S2. Distribution of human metalloproteins with ClinVar pathogenic variants by metal type.** The bars show human metalloenzymes (n = 622) and non-catalytic metalloproteins (n = 378) with at least one ClinVar pathogenic variant and present in the UniProt reference proteome. Each colored segment within a bar represents the proportion of proteins with pathogenic variants that bind a given metal type. A protein is counted once for each metal type it binds.
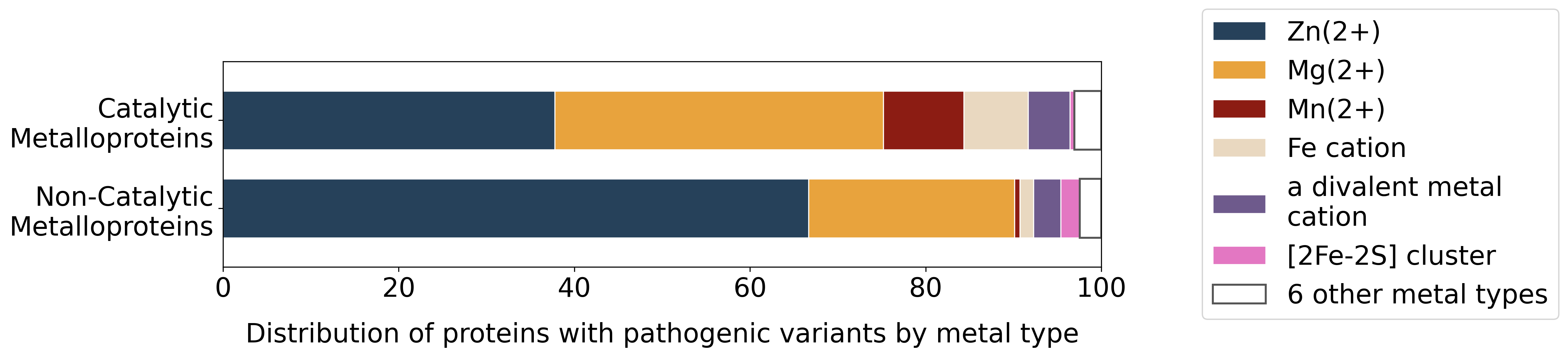

**Table S1. Enrichment of KEGG biological pathway annotation by protein group.** Each group was tested for the proportion of its proteins annotated in at least one KEGG biological pathway, relative to the proportion among all human proteins in the analysis set (8,633/19,576, 44.1%). Expected number of proteins annotated in at least one pathway = n × 0.441, where n is the number of proteins in the group. Fold change = observed count / expected count. A two-sided binomial test was performed for each protein group, with Benjamini–Hochberg correction. Analysis was restricted to proteins with a gene in the KEGG GENES database.

| **Protein group** | **n** | **Observed** | **%** | **Expected** | **Fold change** | ***p*** | ***p*_adj_** | **Result** |
| --- | --- | --- | --- | --- | --- | --- | --- | --- |
| Metalloenzymes | 1,042 | 667 | 64.0 | 459.5 | 1.45 | 4.4 × 10^−38^ | 1.3 × 10^−37^ *** | Enriched |
| Non-catalytic metalloproteins | 747 | 380 | 50.9 | 329.4 | 1.15 | 2.3 × 10^−4^ | 2.3 × 10^−4^ *** | Enriched |
| Other human proteins | 17,787 | 7,586 | 42.6 | 7,844.1 | 0.97 | 9.8 × 10^−5^ | 1.5 × 10^−4^ *** | Depleted |
| All human proteins | 19,576 | 8,633 | 44.1 | — | — | — | — | Reference |

*Enriched or Depleted denotes p_adj_ < 0.05 with fold change above or below 1, respectively. * p_adj_ < 0.05; ** p_adj_ < 0.01; *** p_adj_ < 0.001.*

**Table S2. Enrichment of protein groups across top-level KEGG pathway classes.** Each group was tested for the proportion of its pathway-annotated proteins in a given pathway class, relative to the proportion of all pathway-annotated human proteins in that class. Each protein was counted once per pathway class it appears in; a protein spanning several classes contributes to each, so counts sum to more than the number of annotated proteins. The expected count for a group in a class is the number of that group’s proteins annotated in at least one KEGG pathway, multiplied by the fraction of all pathway-annotated human proteins belonging to that class. Fold change = observed count / expected count. A two-sided binomial test was performed for each protein group–pathway class association, with Benjamini–Hochberg correction. Analysis was restricted to proteins annotated in at least one KEGG pathway.

| **Pathway class** | **Protein group** | **Observed** | **Expected** | **Fold change** | ***p*** | ***p*_adj_** | **Result** |
| --- | --- | --- | --- | --- | --- | --- | --- |
| Cellular Processes | Metalloenzymes | 160 | 181.9 | 0.88 | 0.061 | 0.10 | Not significant |
|  | Non-catalytic metalloproteins | 149 | 103.6 | 1.44 | 4.6 × 10^−7^ | 2.1 × 10^−6^ *** | Enriched |
|  | Other human proteins | 2,045 | 2,068.5 | 0.99 | 0.55 | 0.65 | Not significant |
| Environmental Information Processing | Metalloenzymes | 191 | 221.9 | 0.86 | 0.011 | 0.022 * | Depleted |
|  | Non-catalytic metalloproteins | 136 | 126.4 | 1.08 | 0.30 | 0.39 | Not significant |
|  | Other human proteins | 2,545 | 2,523.7 | 1.01 | 0.61 | 0.65 | Not significant |
| Genetic Information Processing | Metalloenzymes | 96 | 126.1 | 0.76 | 0.0025 | 0.0057 ** | Depleted |
|  | Non-catalytic metalloproteins | 117 | 71.8 | 1.63 | 2.8 × 10^−8^ | 2.5 × 10^−7^ *** | Enriched |
|  | Other human proteins | 1,419 | 1,434.1 | 0.99 | 0.67 | 0.67 | Not significant |
| Human Diseases | Metalloenzymes | 228 | 242.4 | 0.94 | 0.26 | 0.36 | Not significant |
|  | Non-catalytic metalloproteins | 174 | 138.1 | 1.26 | 1.8 × 10^−4^ | 5.4 × 10^−4^ *** | Enriched |
|  | Other human proteins | 2,735 | 2,756.5 | 0.99 | 0.62 | 0.65 | Not significant |
| Metabolism | Metalloenzymes | 341 | 134.0 | 2.54 | 8.5 × 10^−71^ | 1.5 × 10^−69^ *** | Enriched |
|  | Non-catalytic metalloproteins | 52 | 76.4 | 0.68 | 0.0013 | 0.0034 ** | Depleted |
|  | Other human proteins | 1,342 | 1,524.6 | 0.88 | 1.1 × 10^−7^ | 6.9 × 10^−7^ *** | Depleted |
| Organismal Systems | Metalloenzymes | 231 | 279.8 | 0.83 | 1.2 × 10^−4^ | 4.3 × 10^−4^ *** | Depleted |
|  | Non-catalytic metalloproteins | 137 | 159.4 | 0.86 | 0.022 | 0.040 * | Depleted |
|  | Other human proteins | 3,253 | 3,181.8 | 1.02 | 0.099 | 0.15 | Not significant |

*Enriched or Depleted denotes p_adj_ < 0.05 with fold change above or below 1, respectively. * p_adj_ < 0.05; ** p_adj_ < 0.01; *** p_adj_ < 0.001.*

**Table S3. Enrichment of KEGG disease pathway annotation by protein group.** Each group was tested for the proportion of its proteins annotated in at least one KEGG disease pathway, relative to the proportion among all human proteins in the analysis set (3,137/19,576; 16.0%). Expected number of proteins annotated in at least one disease pathway = n x 0.16, where n is the number of proteins in the group. Fold change = observed count / expected count. A two-sided binomial test was performed for each protein group, with Benjamini–Hochberg correction. Analysis was restricted to proteins with a gene in the KEGG GENES database.

| **Protein group** | **n** | **Observed** | **%** | **Expected** | **Fold change** | ***p*** | ***p*_adj_** | **Result** |
| --- | --- | --- | --- | --- | --- | --- | --- | --- |
| Metalloenzymes | 1,042 | 228 | 21.9 | 167.0 | 1.37 | 7.3 × 10^−7^ | 1.1 × 10^−6^ *** | Enriched |
| Non-catalytic metalloproteins | 747 | 174 | 23.3 | 119.7 | 1.45 | 2.5 × 10^−7^ | 7.5 × 10^−7^ *** | Enriched |
| Other human proteins | 17,787 | 2,735 | 15.4 | 2,850.3 | 0.96 | 0.018 | 0.018 * | Depleted |
| All human proteins | 19,576 | 3,137 | 16.0 | — | — | — | — | Reference |

*Enriched or Depleted denotes p_adj_ < 0.05 with fold change above or below 1, respectively. * p_adj_ < 0.05; ** p_adj_ < 0.01; *** p_adj_ < 0.001.*

**Table S4. Enrichment of protein groups across individual KEGG disease pathways.** Each group was tested for the proportion of its proteins in a given disease pathway, relative to that group’s overall proportion among all disease pathway-annotated human proteins. Expected count = number of proteins in that disease pathway multiplied by the group’s overall proportion among disease pathway-annotated human proteins; fold change = observed count / expected count. A two-sided binomial test was performed for each protein group–disease pathway association, with Benjamini–Hochberg correction. Analysis was restricted to proteins annotated in at least one KEGG disease pathway. (.xlsx)

**Table S5. Enrichment of ClinVar pathogenic variant annotation by protein group.** Each group was tested for the proportion of its proteins whose gene carries at least one pathogenic or likely pathogenic ClinVar variant, relative to the rate among all human proteins in this analysis set (56,679/144,748; 39.2%). Expected number of proteins with at least one pathogenic variant = n x 0.392, where n is the number of proteins in the group. Fold change = observed count / expected count. A two-sided binomial test was performed for each protein group, with Benjamini–Hochberg correction. Analysis was restricted to proteins with a gene in ClinVar.

| **Protein group** | **n** | **Annotated** | **%** | **Expected** | **Fold change** | ***p*** | ***p*_adj_** | **Result** |
| --- | --- | --- | --- | --- | --- | --- | --- | --- |
| Metalloenzymes | 1,340 | 622 | — | 524.7 | 1.19 | 7.3 × 10^−8^ | 2.2 × 10^−7^ *** | Enriched |
| Non-catalytic metalloproteins | 908 | 378 | — | 355.5 | 1.06 | 0.13 | 0.20 | Not significant |
| Other human proteins | 142,500 | 55,679 | — | 55,798.8 | 1.00 | 0.52 | 0.52 | Not significant |
| All human proteins | 144,748 | 56,679 | 16.0 | — | — | — | — | Reference |

*Enriched or Depleted denotes p_adj_ < 0.05 with fold change above or below 1, respectively. * p_adj_ < 0.05; ** p_adj_ < 0.01; *** p_adj_ < 0.001.*

**Table S6. Enrichment of protein groups across ClinVar diseases.** Each group was tested for the proportion of its protein–pathogenic variant associations for a given disease, relative to that group’s overall proportion among all protein–pathogenic variant associations. Expected count = number of associations for a given disease multiplied by the group’s overall proportion among protein–pathogenic variant associations; fold change = observed count / expected count. A two-sided binomial test was performed for each protein group–disease association, with Benjamini–Hochberg correction. Analysis was restricted to proteins with at least one ClinVar pathogenic variant. (.xlsx)

**Table S7. Log of AI-assisted disease name consolidation operations.** Every name change made during the AI-assisted consolidation step, one row per operation. *change_type* gives the category assigned to the operation, *original name* and *consolidated name* the disease names before and after, and *reason* the model’s stated justification. Input was the 12,957 disease names remaining after MONDO-based consolidation. Produced with Claude Code v.2.1.116; the prompt is stated in Supplementary Text S1. (.xlsx)

**Table S8. HEAM metalloproteins and their catalytic/non-catalytic classifications.** (.xlsx)

**Supplementary Text S1. Prompt used for AI-assisted disease name consolidation.** Disease name consolidation was performed with Claude Code v2.1.116 in August 2026. The input file contained a list of 12,957 disease names remaining after MONDO-based consolidation. The prompt is stated below.

Create a file called disease_name_changes.tsv in this directory. it will have these sections:
 “change_type”, “original name”, “consolidated name”, and “reason”. Read cleaned_disease_
counts.txt. A) find any disease entries that are actually multiple names and split them into
individual disease names. Log every occurrence of this to disease_name_changes.tsv and state
the reason for splitting. B) find any disease entries that are synonyms of each other and merge
them together. log this in the disease_name_changes.tsv. C) find any disease entries that are
subtypes of an existing disease entry in cleaned_disease_counts.txt. log this in the file. this file
will be read by future scripts to make updated protein-pathogenic variant mapping csv files.
